# ATP allosterically stabilizes Integrin-linked kinase for efficient force generation

**DOI:** 10.1101/2021.03.30.437490

**Authors:** Isabel M. Martin, Michele M. Nava, Sara A. Wickström, Frauke Gräter

## Abstract

Focal adhesions link the actomyosin cytoskeleton to the extracellular matrix regulating cell adhesion, shape, and migration. Adhesions are dynamically assembled and disassembled in response to extrinsic and intrinsic forces, but how the essential adhesion component intergrin-linked kinase (ILK) dynamically responds to mechanical force and what role ATP bound to this pseudokinase plays remains elusive. Here, we apply force-probe molecular dynamics simulations of human ILK:*α*-parvin coupled to traction force microscopy to explore ILK mechanotransducing functions. We identify two key saltbridge-forming arginines within the allosteric, ATP-dependent force-propagation network of ILK. Disrupting this network by mutation impedes parvin binding, focal adhesion stabilization, force generation, and thus migration. Under tension, ATP shifts the balance from rupture of the complex to protein unfolding, indicating that ATP increases the force threshold required for focal adhesion disassembly. Our study proposes a new role of ATP as an obligatory binding partner for structural and mechanical integrity of the pseudokinase ILK, ensuring efficient cellular force generation and migration.

## Introduction

Cells sense and respond to a broad variety of biochemical and mechanical signals from the neighboring cells and the surrounding microenvironment, including the extracellular matrix (ECM). Produced and remodeled by the cells, the ECM serves as a physical scaffold for tissues but also actively guides tissue development and homeostasis by regulating a broad range of cellular processes such as adhesion, migration, growth and differentiation (Hynes, 2002; Legate et al., 2009). ECM assembly and bidirectional cell-ECM signaling is mediated by the integrin family of cellular surface receptors. Integrin-ECM binding leads to recruitment of filamentous (F-) actin to allow force generation though the contractile actomyosin cytoskeleton. As integrins lack enzymatic activity and do not bind actin directly, their function depends on establishing and maintaining large, multiprotein complexes of actin-binding and -regulatory proteins, the focal adhesions (FAs) (Schiller et al., 2011). FAs are crucial for the precise spatio-temporal coordination of integrin signaling. In addition to sensing ECM composition, integrins and FAs sense mechanical cues and transduce them into biochemical signals through a process termed mechanotransduction (Hoffman et al., 2011). These mechanical cues include ECM rigidity, tension and shear. A key event of mechanosensing is modulation of actomyosin contractility and FA dynamics, determined by the balance of protein association and dissociation. According to current models, collectively termed the molecular clutch model, exposing cells to large, rapidly applied forces or plating cells stiff substrates leads to large FAs with slower exchange rates of adapter molecules and longer FA lifetimes (Elosegui-Artola et al., 2018). Thus, precise coordination of the stability of FAs and their actomyosin linkage is required for effective generation of traction stresses during mechanosensing as well as during directed cell migration (Hoffman et al., 2011;Kechagia et al., 2019).

A central regulator of FA dynamics and stability downstream of *β* 1 integrins is integrin-linked kinase (ILK), one of the few essential and evolutionarily conserved components of FAs (Wu, 2004; Legate et al., 2006). ILK is the central component of a tripartite IPP-complex comprising ILK, PINCH (particularly interesting new cysteine-histidine rich protein) and *α*-parvin (Wickström et al., 2010). ILK consists of two distinct domains (Fig. 1 A,B): an N-terminal ankyrin-repeat domain, associated with PINCH (Chiswell et al., 2008) and a C-terminal atypical kinase domain which binds to the calponin-homology (CH2) domain of *α*-parvin (Fukuda et al., 2009). The current notion is that the IPP serves as a signal processing platform by recruiting a variety of proteins. For example, *α*-parvin directly interacts with paxillin (Nikolopoulos and Turner, 2001; Lorenz et al., 2008; Wang et al., 2008), PINCH influences receptor tyrosine kinases (Velyvis et al., 2003), and both parvin and PINCH bind F-actin (Vaynberg et al.,2018; Yang et al., 2021) connecting the IPP to the cytoskeleton. Although ILK was believed to directly associate with *β*-integrin tails (Hannigan et al., 1996) more recent studies show that ILK rather indirectly contacts integrins by binding to kindlin-2 (Montanez et al., 2008; Kadry et al., 2018). As kindlin-2 directly interacts with β-integrin (Calderwood et al.,2013; Harburger et al., 2009; Li et al., 2017; Qadota et al.,2012; Jahed et al., 2019) and binds PIP2 via its PH-domain (Liu et al., 2011, 2012), this interaction recruits the IPP to the cell membrane at sites of FAs. While ILK contains a kinase domain with a typical kinase fold, the catalytic function of ILK has been heavily debated (Hannigan et al., 2005; Legate et al., 2006; Fukuda et al., 2009, 2011; Maydan et al., 2010; Wickstrom et al., 2010; Hannigan et al., 2011) and ILK is now is widely regarded as a pseudokinase. Its obligate interaction partner parvin binds to the putative substrate entry site within the kinase-like domain and disruption of ILK:parvin binding reduces the localization of ILK to FAs (Fukuda et al.,2009). Interestingly, ILK retained its ability to bind ATP in the nucleotide-binding cleft in an unusual binding mode (Fukuda et al., 2009) and recent work has identified a role for ATP binding in actin stress fiber formation and adhesion morphology (Vaynberg et al., 2018). However, the molecular mechanism by which ATP impacts ILK functions remains fairly elusive.

**Figure 1.**
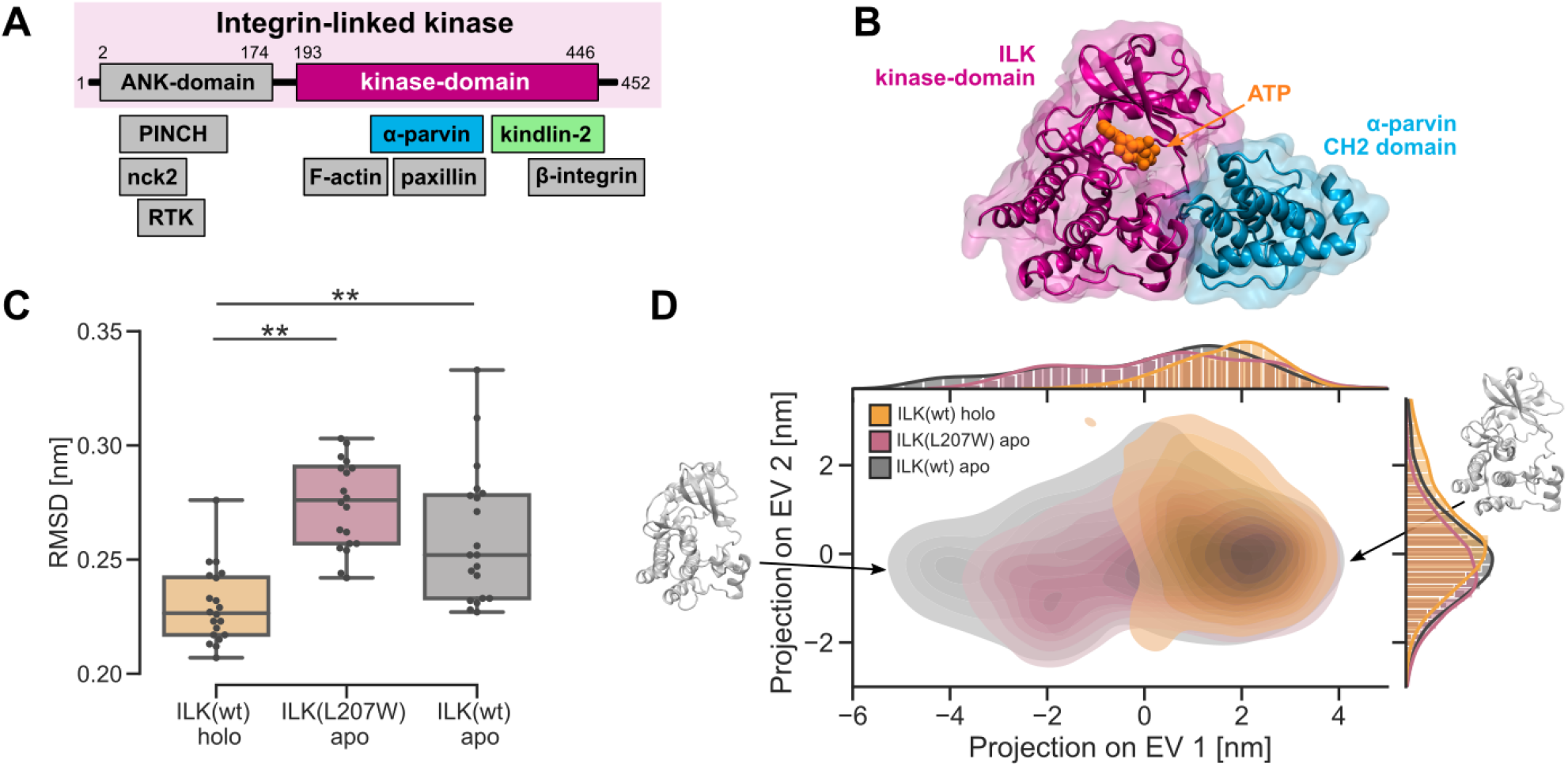
Destabilization of ILK pseudokinase without ATP. (A) Schematic overview of ILK and its associated proteins. (B) The ILK:parvin complex rendered from PDB-code: 3KMW (Fukuda et al., 2009). (C) Trajectory median backbone RMSD of ILK(WT) holo and apo and apo ILK(L207W). n=20 trajectories; ** p = 0.001, one-way ANOVA/ Tukey HSD. (D) PCA of holo and apo ILK(WT) and ILK(L207W). Structures extracted from MD simulations are projected onto PC axes for the first and second PC. Exemplary conformations of apo ILK at two extreme conformations along PC1 are shown.

The importance of the IPP in regulating the integrin-actin linkage is apparent in deletion studies in mice and cells: ILK depletion results in embryonic lethality caused by failure in epiblast polarization and severe defects in F-actin organization at adhesion sites (Sakai et al., 2003). Furthermore, tissuespecific deletion of ILK leads to heart disease (Hannigan et al.,2007) while ILK is also associated with cancer progression (Zheng et al., 2019) and might play a role in ageing (Olmos et al., 2017). At the cellular level, ILK deficiency leads to aberrant remodelling of the actin and microtubule cytoskeletons and decreased force generation, which comprises FA formation, cell migration and ECM remodeling (Friedrich et al., 2004; Fukuda et al., 2003; Sakai et al., 2003; Stanchi et al., 2009; Wickström et al., 2010).

While ILK is clearly indispensable for the integrin-actin connection and efficient force generation through integrins, the precise molecular mechanism of force-induced IPP signalling and the role of ATP therein are unknown. One critical determinant of FA kinetics and thus mechanosensing is thought to be force-sensitive changes in protein conformations. These changes depend on the strength and duration of force application: forces must be transmitted for a period sufficient to induce conformational changes, but large forces also trigger bond breakage or protein dissociation, which will terminate force transmission. Thus, a constant competition between conformational change and bond breakage likely exists at adhesions under high traction stresses (Hoffman et al.,2011; Kechagia et al., 2019). Molecular examples include Focal Adhesion Kinase and talin: the competition between force-induced unfolding and rupture from binding partners depend on parameters such as loading rate and direction, and define their mechanosensing roles (Zhou et al., 2015; Tapia-Rojo et al., 2020; Goult et al., 2018; Mofrad et al., 2004; Rahikainen et al., 2019; Bauer et al., 2019).

Regulated by the activity of the chaperone Hsp90, the stability of ILK is critical for the ability of cells to generate traction forces (Radovanac et al., 2013). This suggests that force-induced unfolding of ILK could play a role for FA signal transduction by promoting disassembly of force-bearing adhesions (Radovanac et al., 2013). To address the mechanosensitivity of ILK and the molecular mechanisms of FA regulation we employ extensive molecular dynamics (MD) simulations of ILK in equilibrium conditions and under mechanical load coupled to biochemical and cellular experiments. Our studies provide evidence for an allosteric strengthening effect of ATP on the ILK:parvin binding and a role for ATP on the cellular force propagation towards *α*-parvin. Furthermore, previously unrecognized ILK saltbridge residues are identified to be critical for ILK:parvin interaction and force transmission. We propose that ILK retained ATP as an obligatory binding partner for structural integrity and efficient force transmission of FAs.

## Results

### ATP structurally stabilizes the ILK pseudokinase domain

To explore the effect of ATP on the ILK pseudokinase dynamics we performed microsecond-scale, all-atom MD simulations with explicit water of wild type (WT) ILK in the holo state (with ATP, PDB-code: 3KMW) and apo state (without ATP, PDB-code: 3KMU) as well as of the ILK(L207W) mutation in the pseudokinase domain that was shown to sterically occlude ATP binding to ILK without affecting the structural integrity of the protein (PDB-code: 6MIB, (Vaynberg et al.,2018)). For each, we simulated the ILK pseudokinase domain in complex with the *α*-parvin CH2-domain to mimic the cellular conditions as closely as possible since parvin is the obligate binding partner of ILK. We find that the overall flexibility of the ILK pseudokinase domain is significantly increased in the apo state as indicated by a higher root-mean-square-deviation (RMSD) from the respective starting structures observed without ATP and similarly in the ILK(L207W) mutation (Fig.1 C). Thus, ATP-binding leads to a decrease in the internal kinase dynamics.

To further characterize the dynamic effect of ATP we examined the differences in the large scale coordinated motions using principle component analysis (PCA, Fig.1 D). We find that in the apo states of both ILK(WT) and ILK(L207W) the pseudokinase explores more of the available conformational space especially along the direction of the most prominent motion (PC1). This motion describes a “wringing”, where the N-lobe of the kinase fold twists around the nucleotide binding pocket on top of the C-lobe (Fig.1 D, movie SV1). Hence, the lack of ATP promotes this wringing motion which contributes to the higher internal flexibility of apo ILK.

### Loss of ILK:ATP-binding destabilizes focal adhesions in particular on rigid substrates

As the simulations showed that ATP is required to stabilize the pseudokinase domain, we sought to functionally test the consequences of this instability in the absence of ATP. To analyze the role of ATP-binding on the ILK:*α*-parvin interaction, we reconstituted ILK-deficient murine fibroblasts (Sakai et al., 2003) with either ILK(WT)-GFP or ILK(L207W)-GFP and performed co-immunoprecipitation assays. Western blot analyses of the immunoprecipitates showed that the L207W mutation had no substantial effect on steady state *α*-parvin binding (Fig.2 A,B) as also reported before (Vaynberg et al.,2018). This was consistent with immunofluorescence analyses of *α*-parvin and ILK localization, which showed a comparable pattern of FA staining of both ILK and parvin in cells expressing ILK(WT) or the ILK(L207W) mutant (Fig.2 C, SFig.2 A).

**Figure 2.**
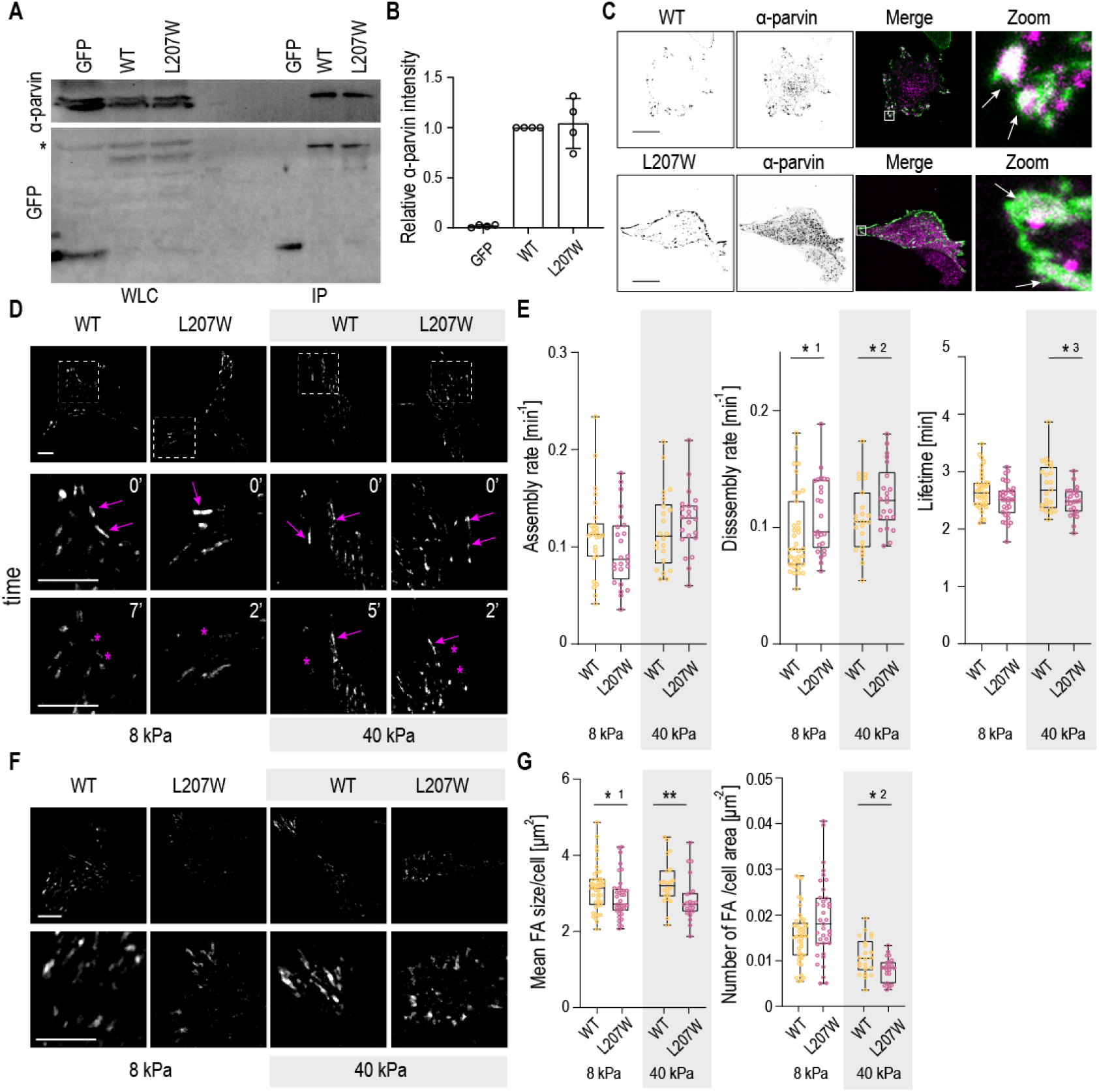
Loss of ILK ATP binding destabilizes focal adhesions in particular on rigid substrates. (A) Representative western blot of GFP pull down experiments from ILK -/- fibroblasts expressing GFP, ILK(WT)-GFP or ILK(L207W)-GFP. *α*-parvin is co-precipitated at similar levels in ILK(WT) and L207W cells. The asterisk marks unspecific antibody binding. (B) Quantification of *α*-parvin to GFP ratio from immunoprecipitation experiments (mean±S.D., n=4 independent experiments). (C) Representative immunofluorescence stainings of *α*-parvin and GFP. *α*-parvin localizes at focal adhesions both in ILK(WT)-GFP and ILK(L207W)-GFP cells. Right panels show a zoom in of the area indicated by the white rectangle. Arrowheads indicate colocalization of *α*-parvin (magenta) with ILK(WT)-GFP and ILK(L207W)-GFP (green). (D) Representative images of ILK(WT)-GFP and ILK(L207W)-GFP cells plated on 8 and 40 kPa substrates and imaged over time to quantify adhesion dynamics. Bottom panels show a zoom in of the area indicated by the dashed square. Arrowheads indicate adhesion growth while asterisks mark adhesion disassembly (Movies SV2, SV3). (E) Quantification of the assembly, disassembly and lifetime of adhesions in ILK(WT)-GFP and ILK(L207W)-GFP cells plated on 8 and 40 kPa substrates (n >5 cells/experiment/condition pooled across 4 independent experiments, *^1^p= 0.0469, *^2^p= 0.0437, *^3^p= 0.0479, Mann-Whitney). (F) Representative images of adhesions in ILK(WT)-GFP and ILK(L207W)-GFP cells on 8 and 40 kPa substrates. Bottom panels show an enlargement. (G) Quantification of the mean focal adhesion size/cell and number of adhesion/cell area in ILK(WT) and ILK(L207W)-GFP cells cultured on 8 and 40 kPa substrates from live imaging. (n >5 cells/condition/experiment pooled across 4 independent experiments, *^1^p= 0.0396, *^2^p= 0.0258, **p= 0.0022, Mann-Whitney). Scale bars 20 *μ*m.

To understand the effect of this mutation on adhesion stability in conditions of low and high traction forces we performed time-lapse imaging of focal adhesion dynamics in ILK(WT)-GFP and ILK(L207W)-GFP cells plated on polyacrylamide gels (PAA) with either low (8 kPa) or high (40 kPa) stiffness. Quantification of adhesion dynamics showed no significant differences in the assembly rates of focal adhesions in WT and L207W mutant, regardless of substrate stiffness. However, the L207W mutant showed faster disassembly rates as well as decreased adhesion lifetimes on both soft and stiff substrates. Interestingly, this difference was more pronounced on the stiff substrates (Fig.2 D,E, Movies SV2, SV3). Consistent with destabilization of adhesions, further analysis of FA morphology revealed overall lower number of FAs per cell area in L207W cells compared to WT (Fig2 F, G), as also reported earlier (Vaynberg et al., 2018). In addition, FAs were slightly smaller in size. Again, these differences were more pronounced on stiff substrates (Fig.2F, G). Decreased FA area was confirmed by paxillin staining on cells plated on crossbow micropatterns to remove cell shape heterogeneity (SFig.2 B) (Théry et al., 2006). Collectively and consistent with our MD simulations, these results indicate that although the inability of the ILK pseudocatalytic domain to bind ATP does not substantially impair the biochemical interaction with *α*-parvin, it destabilizes FAs, in particular under high traction stresses that occur on rigid substrates.

### ATP allosterically influences parvin-binding saltbridges

The pronounced change in the internal collective kinase dynamics and the effects on FA stability suggested that ATP not only affects the residues in its immediate binding pocket but should also exhibit long range effects within the kinase-like domain. We thus next asked which residues and interactions are specifically altered by ATP on a molecular level. The per-residue root-mean-square-fluctuation (RMSF) showed minor differences across a majority of ILK residues between the holo and apo state (SFig.3.1).

To pinpoint residue-residue interactions critical for the ATP induced altered kinase dynamics we performed force distribution analysis (FDA, (Costescu and Gräter, 2013)). Here, the residue-based pairwise forces are calculated and averaged over the trajectory. We measured the force distribution in the holo complex and calculated the differences to the apo structure as a reference to detect ATP-induced allosteric pathways. Fig.3 A shows the network of residue pairs that exhibit changes in interresidue forces larger than a given threshold. Not surprisingly, many residues composing the ATP binding pocket adapt upon ATP binding (Fig.3A, orange arrow).

**Figure 3.**
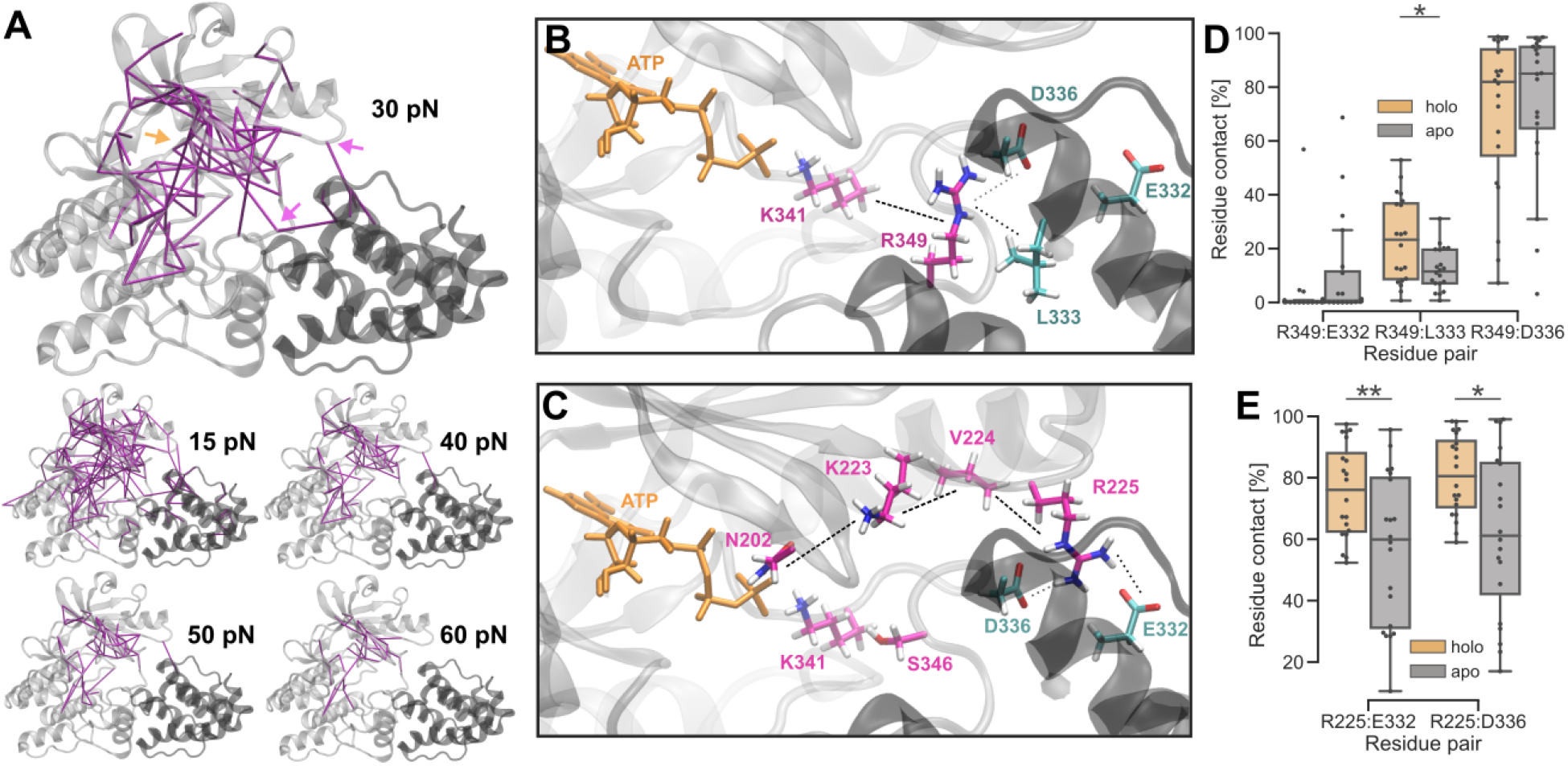
ATP allosterically influences parvin-binding saltbridges. (A) Internal residue-based forces compared between ILK holo and apo with different force thresholds (purple lines), approximate position of the ATP-binding pocket (orange arrow) and saltbridge-forming arginines R225 and R349 (purple arrows). (B,C) Saltbridges between ILK and parvin and proposed force transduction pathway from FDA. (D) Trajectory averaged residue contact probability (threshold 0.35 nm) between ILK R349 and parvin depending on the presence of ATP (n = 20 independent trajectories. *p= 0.017, Mann-Whitney). (E) Residue contact probability between ILK R225 and parvin as in D (n = 20 independent trajectories.**p= 0.0096, *p= 0.013, Mann-Whitney).

ATP also strongly affected residues outside of the binding pocket. Changes in force distribution patterns extended into the kinase N- and C-lobe and strikingly even into parvin. This suggested that ATP has an allosteric effect on the obligatory binding partner of ILK. Particularly important for this intermolecular allostery were two arginine residues: R225 and R349 at the ILK:parvin interface (marked with purple arrows in Fig.3 A). R225 is located on a loop of the N-lobe and R349 belongs to the activation segment of conventional kinases. Structural inspection of these arginines revealed that they form previously unrecognized saltbridges with E332 and D336 in parvin and R349 additionally interacts with L333 of parvin (Fig.3 B,C). Another intermolecular residue pair identified by FDA was D226 of ILK and Q335 of parvin (visible at 15 pN force threshold). The pathway of internal force changes proposed by FDA from ATP to R349 proceeds over the ATP-coordinating K341, corresponding to glycine from the crucial DFG motif of conventional kinases (Fig.3B, SFig.3.2 A) (Fukuda et al., 2009). R225 is found to be allosterically influenced by ATP over a residue cascade involving N202, K223 and V224 (Fig.3 C, SFig.3.2 B). FDA also provided hints towards a pathway involving K341 and S346, however without statistical significance (SFig.3.2 B).

Previous investigations of the ILK:parvin binding interface primarily focused on the parvin binding helix (**α**-G-helix) containing M402 and K403, which mutated to alanine lead to a loss of ILK:parvin binding (Fukuda et al., 2009). To further confirm the importance of the two saltbridge-forming arginines, we performed MD simulations of holo ILK(WT) without parvin. In addition to the parvin-binding helix, the two saltbridge-forming arginines R225 and R349 show a higher flexibility in the absence of parvin (SFig.3.3) which indicates that one of their main functions is to bind parvin. The D336-Q335 residue pair, however, was not visibly perturbed by the lack of parvin and can therefore be regarded as only of minor importance in parvin binding. To further asses the ability of the arginines to form saltbridges with parvin we calculated the saltbridge occupancy/residue contact probability (percentage of total simulation time where the two residues are in contact below a threshold, Fig.3 D, E). The average occupancy of around 80% for the R225-E332, R225-D336 and R349-D336 saltbridges corroborates that those are indeed important functional parvin binding residues. The R349-L333 contact percentage is around 25% on average, rendering this residue pair less important for parvin binding. We confirmed the FDA results on the ATP-dependent ILK:parvin interactions by comparing the saltbridge occupancy/residue contact probability across the interface as observed in the ATP bound state to the apo state (Fig.3 D, E). We found that the absence of ATP significantly lowered the occupancy of the R225-E332 and the R225-D336 saltbridges which suggests an allosteric destabilization of the N-lobe:parvin contacts. The two saltbridges involving R349 did not show statistically significant changes upon ATP removal, however the R349-L333 contact is also significantly reduced. Thus, the lack of ATP does not fully abolish the saltbridges/contacts between ILK and parvin but appear to weaken the interface. Overall, our data suggested that the functional parvin binding interface extends towards the ILK activation loop and especially the N-lobe with R225 and R349, interactions which contribute to the stability of the complex.

### Point mutations in the saltbridge-coordinating residues destabilize *α*-parvin binding and focal adhesions

To understand the functional role of the two saltbridge residues R255 and R349 that we identified by the simulations to mediate the allosteric effect of ATP on ILK:parvin binding, we generated single R255A and R349A as well as a R255A/R349A double mutant, reconstituted ILK -/- cells with these proteins, and performed co-immunoprecipitation assays with *α*-parvin. Both R255A and R349A single mutants showed reduced *α*-parvin binding compared to ILK(WT), whereas strikingly R255A/R349A showed strong reduction of parvin binding (Fig.4 A,B). Immunofluorescence analyses further revealed that although the R255A/R349A double mutant was able to localize to FAs, *α*-parvin failed to efficiently do so and instead displayed a punctate cytoplasmic staining pattern (Fig.4C, SFig.4 A).

**Figure 4.**
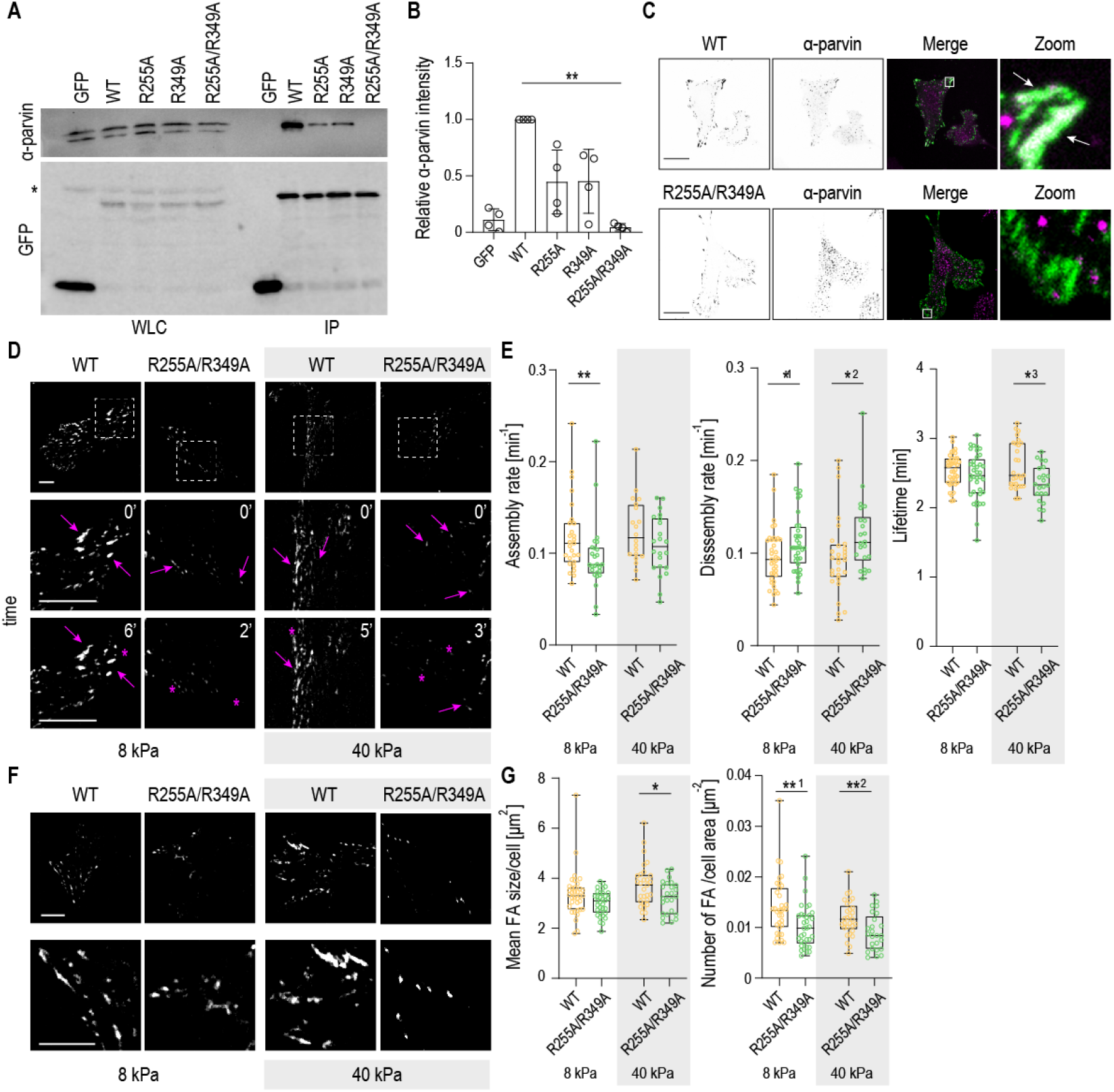
Point mutations in the predicted saltbridge-coordinating residues destabilize *α*-parvin binding and focal adhesions. (A) Representative western blot of GFP pull down experiments from ILK -/- fibroblasts expressing GFP, ILK(WT)-GFP, ILK(R225A)-GFP, ILK(R349A)-GFP and ILK(R225A/R349A)-GFP mutants. R225A-GFP and R349A-GFP show reduced *α*-parvin binding and ILK-(R225A/R349A) complete loss of interaction. The asterisk marks unspecific antibody binding. (B) Quantification of *α*-parvin to GFP ratio from immunoprecipitation experiments (mean ± S.D., n=4 independent experiments, **p= 0.0035, Friedman/Dunn). (C) Representative immunofluorescence images of **α**-parvin and ILK. *α*-parvin co-localizes at focal adhesions with ILK(WT)-GFP cells (arrowheads) whereas no obvious localization of *α*-parvin to adhesions is seen in ILK-(R225A/R349A)-GFP cells. Right panels show zoom in of the area indicated by the white rectangle. (D) Representative images of ILK(WT)-GFP and ILK(R225A/R349A)-GFP cells plated on 8 and 40 kPa substrates and imaged overt time to assess adhesion dynamics. Bottom panels show an enlargement of the area. Arrowheads indicate adhesion growth while asterisks mark adhesion disassembly (Movies SV4, SV5). (E) Quantification of adhesion assembly, disassembly and lifetime in ILK(WT)-GFP and ILK(R225A/R349A)-GFP cells on 8 and 40 kPa substrates (n >5 cells/experiment/condition pooled across 4 independent experiments, *^1^p= 0.0394, *^2^p= 0.0210, *^3^p= 0.0113, **p = 0.0084, Mann-Whitney). (F) Representative images of focal adhesions in ILK(WT)-GFP and ILK(R225A/R349A)-GFP cells on 8 and 40 kPa substrates. Bottom panels show an enlargement of the area indicated by the black square. (G) Quantification of the mean focal adhesion size and adhesion number in ILK(WT)-GFP and ILK(R225A/R349A)-GFP cells on 8 and 40 kPa substrates. ILK(R225A/R349A)-GFP cells (n >5 cells/experiment/condition pooled across 4 independent experiments, *p= 0.0454, **^1^p= 0.0016, *^2^p= 0.0054, Mann-Whitney). Scale bars 20 *μ*m.

To understand the consequences of the saltbridge mutations under high and low traction forces, we performed quantitative analysis of the adhesion dynamics in ILK(WT)-GFP and ILK(R255A/R349A)-GFP cells plated on soft (8 kPa) or stiff (40 kPa) substrates. In contrast to the ATP-binding mutant, ILK(R255A/R349A)-GFP cells showed decreased FA assembly rates on both soft and stiff substrates. In addition, and resembling the ATP-binding mutant, increased disassembly rates and decreased adhesion lifetimes were observed, in particular on stiff substrates (Fig.4 D,E, Movies SV4, SV5).

Consistently, morphological analyses of FAs revealed a significantly smaller size and a lower number of adhesions per cell area in ILK(R255A/R349A) cells compared to WT, with a more pronounced phenotype on 40 kPa stiff gels. Decreased FA area was confirmed by paxillin staining on cells plated on crossbow micropatterns (SFig.4 B). Collectively these results indicate that the disruption of ATP induced allostery, which abolishes **α**-parvin binding, leads to slower adhesion assembly as well as destabilization of FAs, in particular on stiff substrates.

### ATP increases the mechanical stability of the ILK:parvin complex

Having observed destabilization of FAs by disruption of ATP binding in particular on stiff substrates with high traction forces, we asked how ATP affects ILK stability under mechanical load using force-probe MD simulations (Fig.5 A). Forces are transmitted from the ECM/integrins to the IPP and the actomyosin network. The interaction between ILK and integrins occurs via kindlin-2, and although a crystal structure of the ILK:kindlin-2 complex is not available, K423 and I427 of ILK have been experimentally validated as residues interacting with kindlin-2 (Kadry et al., 2018). We determined a larger patch between ILK and kindlin-2 which includes these two residues using homology modelling and guided molecular docking. We defined those residues of ILK that were reproducibly identified to interact with kindlin-2 across the docked ensemble as the kindlin-2 binding patch and as the most probable force-bearing residues on the ILK surface towards the kindlin/integrin anchorage (Fig.5 B). On the other side of the IPP force propagation pathway, *α*-parvin binds actin with its N-terminus over the recently discovered WASP-homology2 domain (Vaynberg et al., 2018) while the N-terminal part of the **α**-parvin CH2 domain is in contact with the LD1 domain of paxillin (Lorenz et al., 2008; Wang et al., 2008). Therefore, the mechanical force transmitted from actin onto the N-terminal part of the CH2 domain would distribute over the residues that are encompassed by paxillin. Those residues were regarded as the pulling patch on parvin (FIg.5 B)

**Figure 5.**
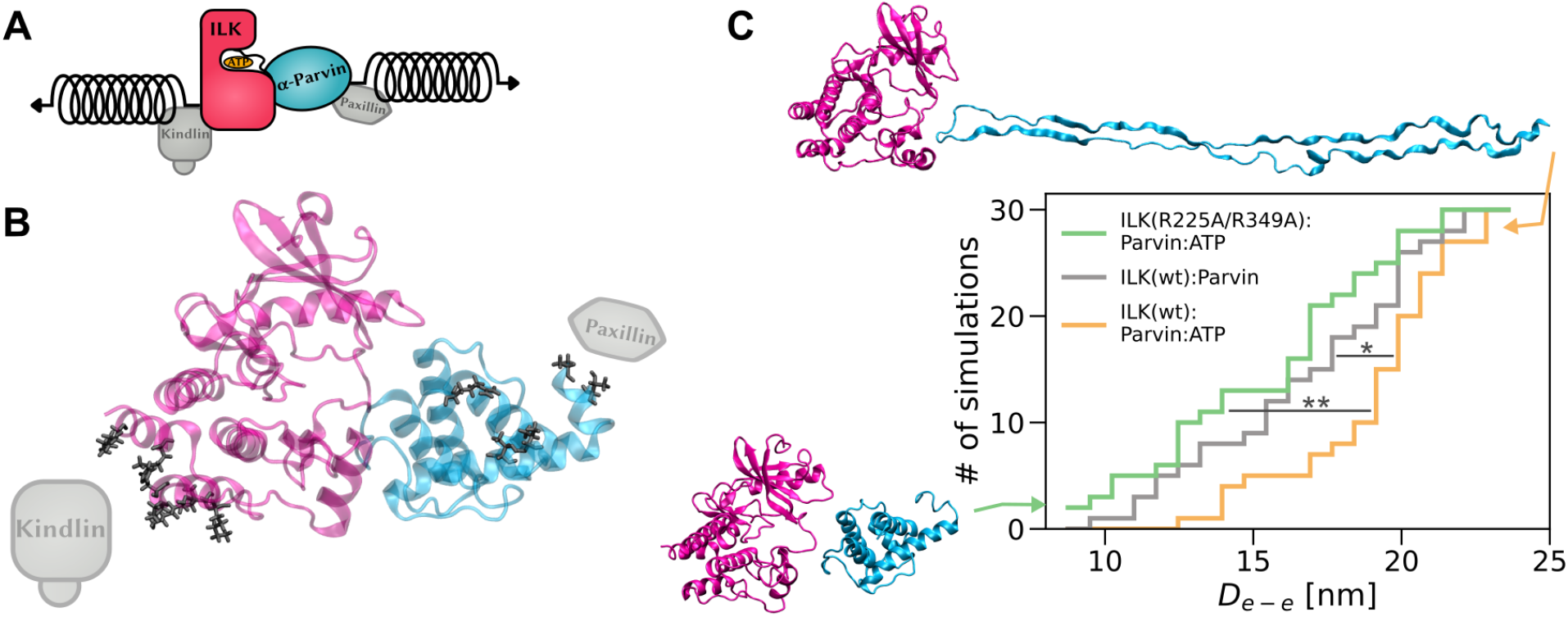
ATP stabilizes ILK:parvin under mechanical tension. (A) Stretching force applied to ILK and parvin in the form of virtual springs at indicated positions. (B) Contact residues (black) between ILK (pink) and a kindlin-2 model (grey) defined as the ILK pulling patch. Contact residues (black) between parvin (cyan) and paxillin-LD1 (grey) defined as the parvin pulling patch. (C) Cumulative number of complex dissociation events as a function of distance between the pulled residue-patches until dissociation (*D_e–e_*) which is defined by an interface area below 0.6 nm^2^ (n = 30 independent trajectories, 10 per velocity, ILK holo vs. ILK apo: p = 0.028, ILK(WT) vs. ILK(R225A/R349A): p = 0.001, one-way ANOVA/Tukey HSD). Example structures of highest and lowest *D_e–e_* before dissociation are shown.

In force-probe MD simulations, we subjected the two sets of force-bearing residues to harmonic pulling potentials moving away from one another with constant pulling velocities of 1 to 0.01 m/s (Fig.5 A,C, SV6 - SV8). Mechanical force induced both ILK:parvin dissociation, as measured by the changes in interface area, and ILK or parvin unfolding, reflected by an increase in the end-to-end distance of the complex prior to dissociation. This general unfolding and dissociation mechanism was similar across different pulling velocities (SFig.5). To quantify and compare the force-induced dynamics, we defined complex dissociation to occur at the time at which the interface area drops below 0.6 nm^2^. Dissociation was observed at various unfolding stages, after only minor parts of the protein complex had straightened and unfolded (end-to-end distance 9 nm) all the way to incidences at which major parts of parvin were already unfolded (end-to-end distance 28 nm). Interestingly, the mechanical stability of the ILK:parvin interface relative to the individual proteins decreases upon depletion of ATP: across pulling velocities, the ATP-bound ILK:parvin complex dissociates at higher extensions, i.e. only after more extensive parvin unfolding, than the apo state. The preference of dissociation over unfolding seen for the wildtype apo complex was also observed in the ATP-bound double saltbridge mutant and was indistinguishable from the wildtype apo complex. These findings indicate a weaker ILK:parvin interface in the absence of ATP which corroborates the notion of ATP as an allosteric regulator of ILK:parvin (Fig.3) and suggests a mechanistic explanation of the observed destabilization of focal adhesions on stiff substrates (Fig.4).

### Loss of ATP- and **α**-parvin binding prevents traction force generation and actomyosin contractility

As the simulations indicated that ATP stabilizes the ILK:parvin complex under mechanical load, we predicted that destabilization of the complex would prevent build up of traction stresses within FAs. To test this, we performed traction force microscopy. Interestingly, both L207W and R255A/R349A showed lower mean traction stresses compared to WT cells (Fig.6 A-D). To assess whether the reduced traction stresses are associated with impaired actomyosin contractility, we analyzed F-actin and myosin II activity by phalloidin and phospho-Myosin Light Chain 2 (pMLC2) stainings. To this end, we plated cells on crossbow-shaped micropatterns to polarize the cells and to accurately quantify changes in myosin II activity across the non-adhesive edges (Théry et al., 2006). WT cells featured thick and mature ventral stress fibers and high pMLC2 (Fig.6 F). Quantification of the immunostainings showed substantial reduction in actin stress fibers and myosin II activity for both the L207W and the R255A/R349A mutants. Collectively, these results show that ATP binding and its allosteric saltbridge interactions are required for the generation of traction stresses and actomyosin contractility.

**Figure 6.**
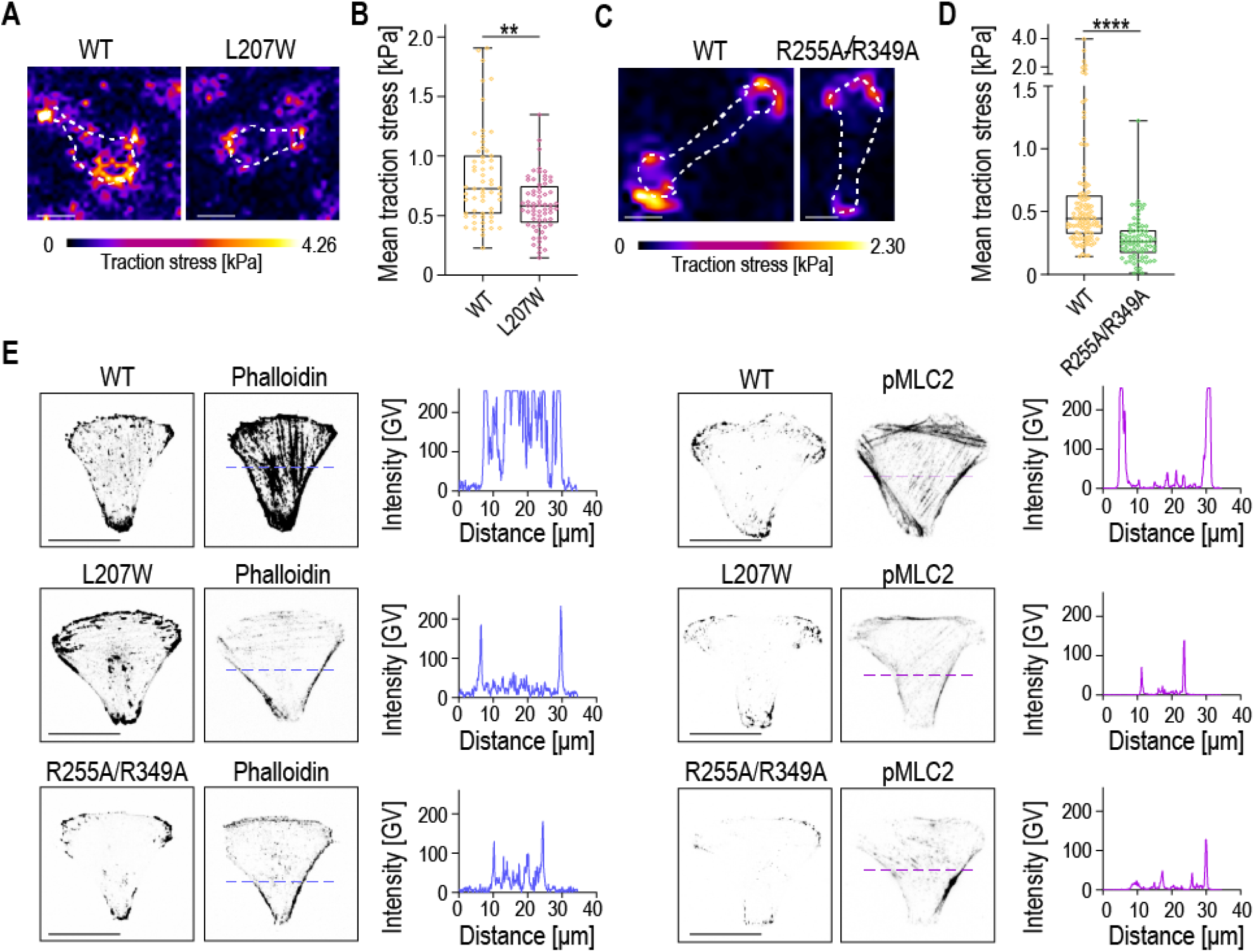
Impaired ATP- and **α**-parvin binding prevent traction force generation and actomyosin contractility. (A) Representative images and traction force heat maps of ILK(WT)-GFP and ILK(L207W)-GFP cells. (B) Quantification of mean traction stresses of ILK(WT) and ILK(L207W) cells (n >10 cells/condition/experiment cells pooled across 4 independent experiments, **p = 0.0047, Kolmogorov-Smirnov). (C) Representative images and traction force heat maps of ILK(WT)-GFP and ILK(R255A/R349A)-GFP cells plated on 8 kPa PAA gels. (D) Quantification of mean traction stresses of ILK(WT)-GFP and ILK(R255A/R349A)-GFP cells (n >10 cells/condition/experiment cells pooled across 4 independent experiments, ****p= 0.0001, Kolmogorov-Smirnov). (E) Representative images of ILK(WT)-GFP, ILK(L207W)-GFP ILK(R255A/R349A)-GFP cells stained with phalloidin and pMLC2 on crossbow micropatterns and corresponding line scan quantifications of phalloidin (in blue) and pMLC2 (magenta) fluorescence intensity. GV = gray values. Scale bars 20*μ*m.

### Loss of ATP- and *α*-parvin binding impairs cell migration

We predicted that the inability of the ILK mutants to generate traction stresses which stabilize FAs and promotes actomyosin contractility would impair cell migration that relies on these processes (Ridley et al., 2003). To test this, we performed cell migration assays and analysed migration trajectories of single cells using live imaging. As expected, the ILK-/- cells showed reduced migration compared to cells where ILK expression was restored by expressing ILK(WT)-GFP (Fig.7 A, B, SV9, SV10). Interestingly, ILK(R255A/R349A)-GFP cells showed a migration defect with migration rates smaller than ILK(WT) cells and comparable to ILK-/- cells (Fig.7 A,B, SV11, SV12). Collectively, these data indicate that destabilization of the ILK:parvin interface prevents build up of traction forces and actomyosin contractility and thus impairs the migratory capacity of cells.

**Figure 7.**
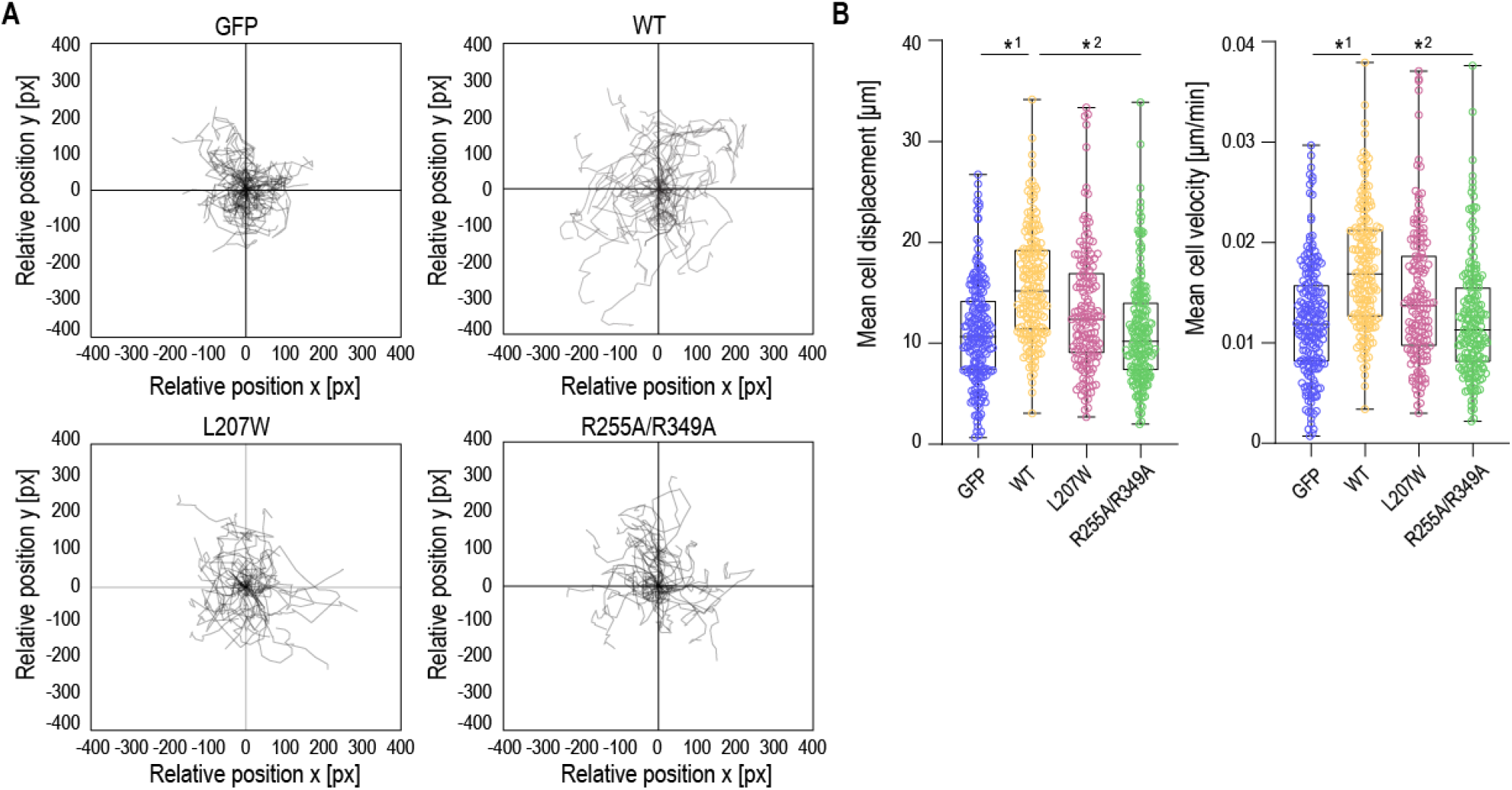
Point mutation in the ATP pseudocatalytic domain and point mutations in the saltbridge-coordinating residues lead to migration defect. (A) Representative trajectory plots showing ILK -/-GFP, ILK(WT)-GFP, ILK(L207W)-GFP and ILK(R255A/R349A)-GFP cell trajectories during 12 h acquisition. (n >30 cell tracks/condition were set to a common origin (intersection of x and y axes). (Movies SV9 - SV12). (B) Quantitative analysis of average distance (*μ*m) and velocity (*μ*m/min) of ILK -/- GFP, ILK(WT)-GFP, ILK(L207W)-GFP and ILK(R255A/R349A)-GFP cells. (n >35 cell/condition/experiment pooled across 3 independent experiments, *^1^p= 0.0114, *^2^p= 0.0269 Friedman/Dunn).

## Discussion

In this study, we employ a combined computational and experimental approach to understand how the pseudokinase ILK integrates and transduces biochemical and mechanical signals. Despite being a pseudokinase, ILK still binds ATP, and we asked if and how ATP plays a role in ILK molecular interactions, function and mechanotransduction. While pseudokinases regulate diverse cellular processes despite lacking the ability to phosphorylate substrates, a significant portion of them bind ATP with largely unknown functions. For other pseudokinases, such as STRAD*α*, ATP was shown to conformationally alter the pseudokinase to maintain a closed conformation for efficient target binding and downstream signaling (Zeqiraj et al., 2009). For ILK, first hints towards a function of ATP were provided by the observation that F-actin bundling is sensitized by ATP bound to ILK (Vaynberg et al., 2018). However, the exact molecular mechanism of ATP function, in particular in the context of mechanical force, has remained elusive. We here put forward a multifaceted role of ATP for ILK function. We find ATP to enhance the structural integrity of ILK, to allosterically impact the ILK:parvin interaction, and to enhance the resistance of the IPP complex to force.

In previous studies, purified ILK was shown to form highly insoluble aggregates in the absence of **α**-parvin (Fukuda et al.,2009) indicating that it likely co-evolved with **α**-parvin as an obligatory binding partner. The inherent instability of ILK is further underlined by the requirement for the heat shock protein Hsp90 for a stable ILK:parvin interaction and force generation on the ECM (Radovanac et al., 2013). Thus, since ATP shows only little effect on the overall ILK structure (Fukuda et al., 2009) our data indicates that ATP instead alters the dynamics and stability of ILK and the ILK:parvin complex proposing ATP as a secondary binding partner for structural integrity.

ATP-binding deficient ILK(L207W) was described as a mutant that sterically occludes the ATP binding site without affecting the structural integrity (Vaynberg et al., 2018). The overall structural integrity is indeed maintained, as shown by the ability of this mutant to still bind parvin ((Vaynberg et al., 2018), this study). However, on an atomistic level the kinase-like domain is more flexible without ATP and displays a more pronounced inter-lobe wringing motion comparable to apo ILK(WT). On the cellular level, loss of ATP in the ILK pseudokinase leads to destabilization of FAs resulting in decreased FA number. This is especially pronounced on stiff substrates where high traction forces are generated.

Our simulation data further provides evidence for internal force propagation pathways from ATP that allosterically influence previously unacknowledged parvin-binding saltbridges (R225 and R349). Indeed, we show that those residues confer parvin binding, herein cooperating with the previously analyzed M402/K403 in the C-lobe of ILK (Fukuda et al., 2009). Interestingly, while failing to recruit parvin to FAs due to impaired binding, the double saltbridge mutant (R349A/R225A) itself still localizes correctly to FAs, contrasting the M402A/K403A mutant. In this way, the double saltbridge mutant phenotypically resembles the ILK mutant deficient in ATP-binding. In contrast to the L207W mutant, another ATP-binding defective mutant (N200A/K341A) failed to localize correctly to FAs (Fukuda et al., 2009). Our force distribution anaylsis indicates that the reason for this might be the importance of K341 for the allosteric signal transduction to the parvin-binding saltbridges. Our data thus puts forward a model that features two distinct parvin binding pathways within ILK: one ATP-dependent pathway across the kinase N-lobe and activation segment which governs the two saltbridge-forming arginines and the previously established C-lobe pathway which is ATP-independent. Intriguingly, only the ATP-independent mode is required for localizing ILK into FAs, whereas ATP-dependent binding is required for adhesion stability.

Beyond the stabilizing effect of ATP on the pseudokinase in equilibrium conditions, we also observe that ATP conveys mechanical stabilization of the ILK:parvin complex under force. The molecular basis for this mechanical stabilization in our simulations is based on the assumption that forces in FAs are transmitted from integrins via kindlin to the IPP ultimately reaching F-actin and vice versa. This is reflected in the choice of residue-patches for the force-probe MD. Using a combined molecular modelling and docking approach for ILK:kindlin, we identify kindlin-binding residues and are able to robustly simulate the resultant pulling direction. A more comprehensive view onto mechanotransduction across the kindlin-ILK axis, potentially also involving force propagation via the membrane as for FAK (Goñi et al., 2014; Zhou et al., 2015), can only be obtained once an experimental kindlin:ILK structure becomes available. Another plausible mechanotransduction pathway involves the N-terminal ankyrin-repeat domains of ILK (Chiswell et al., 2008). Given the critical role of the pulling direction on protein mechanical response (Carrion-Vazquez et al., 2003), this additional force application route might further alter the competition between unfolding and rupture of the IPP complex and the role of ATP therein.

As evident by the indistinguishable mechanical instability of apo ILK and the saltbridge mutant that still binds ATP, we hypothesize that the stabilizing effect of ATP on ILK:parvin under mechanical force is at least partly mediated by the allosterically influenced saltbridges. Under the assumption that an intact ILK:parvin complex is a prerequisite for cellular force generation we therefore predicted that the double saltbridge mutant is impaired in establishing cellular forces due to its apparently weaker interface under mechanical load. Experimental quantification of cell-ECM traction stresses showed that indeed disruption of both ATP- and parvin-binding in ILK leads to reduced force generation and cell migration, supporting the notion that stability of ILK:parvin interaction determines the ability of FAs to bear loads. Thus, we propose that ILK functions as a pseudo-mechanokinase, where ATP binding serves a mechanosensory role. We generally observe a slightly milder phenotype for the ATP-binding mutant compared to the double-saltbridge mutant, which might be explained by fact that parvin is not recruited to FAs in the double-saltbridge mutant. Since the IPP was shown to form in cells in suspension, parvin binding likely precedes recruitment of IPP to FAs (Zhang et al., 2002). Therefore our results of reduced FA assembly rates in the double-saltbridge mutant that strongly disrupts parvin binding might be a hint towards a function of parvin in the IPP-driven FA assembly. In contrast, FA disassembly, especially on stiff substrates, is more severely perturbed in both mutants which might indicate that both parvin and ATP regulate IPP stability under mechanical force. Hence, ATP might be regarded as one puzzle piece that keeps the ILK:parvin complex intact to precisely regulate the point of focal adhesion disruption, determined by the amount of force build-up, which ultimately leads to FA disassembly for efficient rear end retraction.

In summary, this study sheds light onto the stabilizing function of ATP on the ILK pseduokinase leading to efficient traction force build-up, FA stabilization and efficient migration. Apart from further advancing the knowledge on ILK-mediated intergrin-actin communication and the dynamics of FAs, our study contributes to the understanding of pseudokinase evolution and function. Pseudokinases have emerged as useful models to study the non-catalytic properties of the kinase fold in isolation (Jacobsen and Murphy, 2017; Kung and Jura, 2016). With several pseudokinases currently under investigation as therapeutic targets (Byrne et al., 2017; Kung and Jura, 2019) further understanding the function of ILK pseduokinase might culminate in therapeutic implementations to specifically target ILK not as a kinase, but a mechanotransducer. Mechanotransducing (pseudo)kinases are uniquely suited to integrate mechanical into other signalling pathways, and more members of this intriguing class of proteins likely remain to be uncovered.

## Methods

### Equilibrium molecular dynamics simulations

The crystal structures of the human ILK(WT) kinase domain in complex with the CH2-domain of *α*-parvin either with ATP (Protein Data Bank (PDB)-code 3KMW) or without ATP (3KMU, (Fukuda et al., 2009)) were used. To achieve starting structures without parvin, those coordinates were deleted. The PDB-code 6MIB (Vaynberg et al., 2018) was used for the simulations with the ILK(L207W) mutation. Point mutations for ILK(R225/R349A) were generated using pymol. All MD simulations were carried out using GROMACS 2018.1 (Abraham et al., 2015). The Amber99sb*-ILDNP force-field (Best and Hummer, 2009; Aliev et al., 2014) and TIP3-water model (Jorgensen et al., 1983) were used. The starting configurations were solvated in the center of a dodecahedron box with (at least) 3 nm between each periodic image. Sodium and chloride ions corresponding to a physiological concentration of 100 mM were added resulting in a system with overall zero charge. An energy minimization was performed with the steepest descent method, followed by 500 ps in the NVT and 500 ps in the NPT ensemble with harmonic constraints on all protein atoms with a force constant of 1000 kJ mol^−1^nm^−1^ to equilibrate water and ions. The production runs were carried out in the NPT ensemble without constraints on heavy atoms. All bonds between hydrogens and protein heavy atoms were constrained using the LINCS algorithm (Hess, 2008). Therefore, a timestep of 2 fs could be used. The temperature was kept constant at T = 300 K using the velocity rescaling thermostat (Bussi et al., 2007) with a coupling time of 0.1 ps. The two temperature coupling groups were (1) all protein and ATP atoms and (2) all water and ions. The pressure was kept constant at 1 bar using the isotropic Parrinello-Rhaman barostat (Parrinello and Rahman, 1981) with a coupling time of 2 ps and a compressibility of 4.5 ×10^−5^ bar^−1^. The neighbors list was updated every 10 fs with the Verlet scheme. A cutoff of 1.0 nm was used for all non-bonded interactions and long-range electrostatic interactions were treated using the Particle mesh Ewald method (PME, (Darden et al., 1993)) with a grid spacing of 0.16 nm with cubic interpolation. If not stated otherwise the systems were simulated for 500 ns. For the analysis the first 20 ns were neglected as equilibration period, as inspected from the protein backbone root-mean square deviation (RMSD). 20 individual production runs were carried out, with random velocities and their own equilibration phases, totaling to a total simulation time used for the analysis of 9.6 μs per condition.

### Guided molecular docking

To determine a patch of residues on ILK that is most likely in contact with kindlin-2 we employed a guided molecular docking approach. One available crystal structure of the nearly complete mouse kindlin-2 is missing only two loops within the F1 domain and the PH-domain (PDB: 5XPY, (Li et al., 2017)). Unfortunately, the ILK-binding helix (PDB: 2MSU, (Fukuda et al., 2014)) is located at the edge of the PH domain and is only partially included in the nearly full structure. Therefore, we combined all available partial structures of kindlin-2 (ILK-binding helix: 2MSU; free PH-domain: 4F7H and 2LKO (Liu et al.,2011, 2012); N-terminus: 2LGX) and aligned them with the main body of kindlin-2 (PDB: 5XPY) using UCSF-Chimera (Pettersen et al., 2004). This structural alignment was performed according to an underlying sequence alignment with the human kindlin-2 generated with T-coffee (DiTommaso et al., 2011). Since the structures of the free PH-domains do not overlap in sequence with the main kindlin body, they were placed near the artificially inserted loop in the F2 domain of the main body structure in such a way that the residues that follow in the sequence have a minimal distance to each other. In doing so six rotations of the PH-domains were generated. For the missing loop within the F1 domain there is no crystal structure, but it is far apart from the ILK binding site and thus we regard it as neglectable for the ILK binding. The final full structures were generated by a homology modeling using MODELLER (Webb and Sali, 2016) based on the positions of the placed fragments. For each of the six conformations, four models were generated. Eleven models in which the F1-loop was not threaded through the protein were selected for further analysis. Those were subjected to a short 20 ns to 30 ns MD simulation (see above for simulation parameters). A cluster analysis over the whole trajectory determined the most populated conformations, which were selected for the guided molecular docking with Haddock2.2 (van Zundert et al., 2016). For ILK, the most populated structure from the 100 ns MD simulation of PDB-code 3KMW was used. The experimentally validated residues of ILK that are in direct contact with kindlin-2 (K423, I427, (Kadry et al., 2018)) were chosen as the actively participating residues. The residues of kindlin-2 (L353, E354, L357, E358) that were shown to interact with ILK (Fukuda et al., 2014) were set as active residues as well. Passive residues were defined automatically around the active residues. Seven docking protocols were successful and used for further analysis.

### Determination of pulling sites

The ILK:kindlin-2 docking poses were used to determine the most physiological cluster of residues for mechanical perturbation. From the 361 highest scoring docking poses we determined the residues in ILK that are in contact with kindlin-2 within a cutoff of 0.35 nm. The residues that occur in most of the docking poses are: P419, H420, K423, I427, K435, M441, K448. These were then taken as the patch of residues for force-probe MD.

For parvin, we adopted a similar approach. From the structures of the parvin-CH2 domain in complex with the paxillin LD1 domain (PDB-codes: 4EDN (Lorenz et al., 2008) and 2K2R (Wang et al., 2008)) the residues on parvin that are interacting with paxillin within the same threshold of 0.35 nm were determined. This resulted in the residues A249, T252, V264, T267, R369 as the patch of residues on parvin for pulling.

### Force-probe molecular dynamics simulations

We used force-probe MD to simulate the effects of mechanical perturbation on the system. The end-conformations of the equilibrium simulations were placed in the center of a rectangular box of 30 Å ×15 Å ×15 Å and rotated so that the axis between the putative pulling points is in line with the x-axis of the box. The system was energy minimized, followed by short simulations in the NVT and NPT ensemble as described above. For the production run the respective residue patches were subjected to harmonic spring potentials with a force constant of 50 kJmol^−^¼m^−1^ moving in opposite directions with a constant velocity (v= 1 ms^−1^ to 0.01 ms^−1^). 10 simulations were performed per velocity and condition. The simulations were stopped upon dissociation of ILK:parvin.

### MD analysis and visualization

The computational analysis was performed with GROMACS tools and post-processed with Python 3. If not stated otherwise, the first 20 ns of each equilibrium simulation were excluded from the analysis. The RMSD of the backbone atoms was calculated in relation to the first frame of the production run. Further, the two starting structures of the holo and apo complex were found to be remarkably similar to each other (Fukuda et al., 2009). Thus, the differences in RMSD do not reflect differences in the overall structure of the ILK pseudokinase domain but rather show that ATP influences the dynamical behavior.

The saltbridge occupancy/residue contact probabilities were calculated by determining the number of frames of the whole trajectory with a residue-residue distance below a threshold of 0.35 nm.

To determine the interface area between ILK and parvin, we calculated the solvent accessible surface area (SASA, (Eisenhaber et al., 1995)) for both proteins alone and for the complex. The interface area can be calculated according to equation 1:

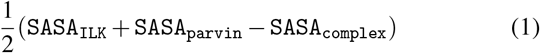

We define a complex dissociation event if the interface area is below 0.6 nm. For visualization of protein structures we used visual molecular dynamics (VMD; (Humphrey et al., 1996)). For the generation of the final figures we used Inkscape.

### Principle component analysis

We performed principle component analysis (PCA, reviewed in (David and Jacobs, 2014)) with GROMACS utilities covar and anaeig to identify the major correlated structural motions. The first 20 ns of each equilibrium MD trajectory were neglected and the PCA was conducted on the C-**α** atoms of the cumulative trajectories of the ILK holo und apo state or ILK(L207W). Rotational and translational motions were removed by superimposing the structures along the trajectory onto the invariant core. A covariance matrix from the C-**α** positions was generated. The eigenvectors that describe the direction of motion, are generated by diagonalization of the covariance matrix. The corresponding eigenvalue describes the magnitude (energetic contribution) of that component to the motion. Projection of the trajectory on the eigenvectors shows the motions of the protein along this mode of motion. All trajectories were projected onto the eigenvectors generated by the apo state. The top two eigenvectors were considered to construct the two-dimensional histogram.

### Force distribution analysis

We used force distribution analysis (FDA (Costescu and Gräter, 2013)) implemented into GROMACS to calculate the changes in internal forces between the ILK holo and apo states. In FDA the pairwise forces between atom pair i and j are calculated considering all interaction types between protein atoms. These time-averaged pairwise forces can be non-zero even in equilibrium simulations. For a residue based analysis the inter-residue forces were calculated according to equation 2:

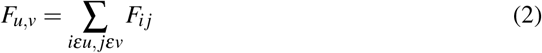

where i is an atom of residue u and j is an atom of residue v, with u and v being different. The FDA was performed on each individual trajectory per condition and the resulting average forces of the apo state were subtracted from those of the holo state. The networks shown are connected graphs of at least 3 residues with force differences above a given threshold. Each interaction in the graphs shows a statistically significant change in internal forces (Mann-Whitney test, p <0.05).

### Plasmid constructs

Full length mouse ILK cDNA was cloned into the EcoRI site of pEGFP-N1 plasmid (Clontech). The R255A, R349A, and L207W mutations were generated by performing site-directed mutagenesis PCR using Quik Change II Mutagenesis Kit (Agilent) with the following primers:

R255A forward: 5’-CCTTGTACTCCAGTCTG CAACCTTCAGCACCTTCA-3’ and
R255A reverse 5’-TGAAGGTGCTGAAGGTTGC AGACTGGAGTACAAGG-3’;
R349A forward 5’-AGGCGCATACATGGCCC CAGGGCACTGG-3’ and
R349A reverse 5’-CCAGTGCCCTGGGGCC ATGTATGCGCCT-3’;
L207W forward: 5’-CCTGCCAGCGGCCTTTCC ACCACTCTCCAGAATGATTCTCA-3’ and
L207W reverse 5’-TGAGAATCATTCTGGAGAG TGGTGGAAAGGCCGCTGGCAGG-3’.

### Cell culture and transfection

Immortalized ILK-/-mouse fibroblasts were obtained as described (Radovanac et al., 2013) and cultured in Dulbecco’s MEM, 10% FBS (Gibco) in 5% CO_2_ at 37 °C. Transient transfections were performed using Lipofectamine 3000 reagent (Invitrogen) according to the manufacturer’s instructions. After 24 h of transfection cells were subjected to specific experimental analyses.

### Immunoprecipitation

Transfected cells grown on polystyrene dishes were rinsed in phosphate-buffered saline (PBS), suspended in lysis buffer (50 mM Tris–HCl buffer (pH 8.0) containing 150 mM NaCl, 1% Triton-X 100, 0.05% sodium deoxycholate, 10 mM EDTA, protease, and phosphatase inhibitors (Roche)), and cleared by centrifugation. GFP immunoprecipitation was performed using Miltenyi Biotec MultiMACS GFP Isolation Kit (Mylteni Biotec 130-091-125) according to the manufacturer’s protocol. The immunocomplexes were eluted in Laemmli sample buffer by boiling and analyzed by western blotting.

### Western Blotting

Lysates were reduced in Laemmli sample buffer at 95°C, separated by PAGE in the presence of SDS, transferred onto PVDF membranes and subjected to western blot analyses using the standard protocols. The following antibodies were used: anti-*α* parvin (Cell Signalling; 4026; 1:2500), anti-GFP (Invitrogen; A1112; 1:2500) and anti-rabbit HRP (Bio-Rad Laboratories). Western blots were quantified using densitometry (ImageJ software) (Schindelin et al., 2012) from four independent experiments.

### Generation of soft and stiff substrates

8 and 40 kPa polyacrylamide (PAA) gels (7.5 % acrylamide/0.25% bis-acrylamide) were cast on 20 x 20 glass coverslips after which fibronectin was chemically crosslinked on gels using Sulfo-SANPAH (Pierce). Gel stiffness was measured as described (Pelham and Wang, 1997). For traction force microscope gels were manufactured as above by additioning 0.2 *μ*m fluorescent beads (1:125; Polysciences). Gels were washed in 70% Ethanol and rinsed extensively in PBS after which transfected cells were plated, allowed to adhere and live imaged for focal adhesion dynamic analysis or used for traction force microscopy experiments.

### Focal adhesion dynamics

Transfected cells on 8 and 40 kPa polyacrylamide (PAA) gels were imaged using a Zeiss Axiovert inverted microscope coupled to a CSUX1 spinning-disc device (Yokogawa) equipped with a 488 nm laser, sCMOS camera (Hamamatsu) and an environment chamber. Imaging was performed with a 100×oil immersion objective (Zeiss) at 37 °C, 5% CO_2_. Images were acquired at a frame rate of 1 frame/30 sec for 30 mins. To quantify adhesion dynamics, time-lapse movies were preprocessed using Fiji (Schindelin et al., 2012) by subtracting the background, denoising and linear contrast enhancement. Image sequences were then submitted to Focal Adhesion Server Analysis FAAS (Berginski and Gomez, 2013). Assembly and disassembly events per adhesion per cell were collected based on R^2^ values greater or equal 0.8 (Berginski and Gomez,2013).

### Traction Force Microscopy

Traction force microscopy was performed essentially as described (Dembo and Wang, 1999). Transfected cells were cultured on 8 kPa PAA. Imaging was performed using a spinning disc microscope described above with a 40x glycerol objective at 37°C with 5% CO_2_. Cells were imaged prior and after the addition of 10X trypsin to detach cells and obtain bead displacement images. Calculation of traction forces was performed using particle imaging velocimetry (PIV) and Fourier transform traction cytometry (FTTC) with regularization (10^−9^) using Fiji (Schindelin et al., 2012) as described previously (Tseng et al., 2011). Traction forces were reconstructed at a grid spacing of 5 *μ*m and total cellular force was calculated from the average of traction magnitudes. At least 30 cells/condition were analysed.

### Migration assay and cell tracking

Transfected cells (5000 cells/well) were plated on 4-well chamber slides (Thermo Scientific^™^ Nunc^™^ Lab-Tek^™^ II Chamber Slide^™^) and imaged by using differential interference contrast (DIC) optics with a Zeiss Axiovert inverted microscope coupled to a CSUX1 spinning-disc device (Yokogawa) equipped with a 488 nm laser, sCMOS camera (Hamamatsu) and an environment chamber. Imaging was performed with a 25× oil immersion objective (Zeiss) at 37 °C, 5% CO_2_. Images were acquired at a frame rate of 1 frame/30 min for 12 _%_hrs. To quantify individual cell trajectories, cells were manually tracked by using the Fiji plugin Trackmate (Tinevez et al.,2017). The mean distance and cell velocity were calculated by averaging the displacement (in *μ*) and the velocity (*μ*m/min) between consecutive frames.

### Micropattern fabrication

Micropatterned adhesive surfaces were generated using the PRIMO optical module (Alveole, France) controlled by the Leonardo plugin (V3.3, Alveóle) mounted on a Nikon TI-E inverted microscope (Nikon Instruments) equipped with a Super Plan FLuor 20x ELWD lens (Nikon) lens and a DMD-based UV (375 nm). Crossbow-shaped micropatterns (length 35 *μ*m, width 17.5 *μ*m, radius 17.5 *μ*m) were projected onto plasma-cleaned (Corona Treater, ETP), PLLgPEG-passivated (0.1 mg/ml PLL-g-PEG (PLL (20)-g [3.5]- PEG (2), SuSoS) 35mm glass-bottom dishes (Ibidi). Patterned areas were then washed multiple times with PBS and conjugated with a uniform coating of 10 μg/ml fibronectin for 1h at 37°C. The substrates were then washed with PBS, prior to seeding 10000 transfected fibroblast cells onto each 35mm dish. Cells were allowed to adhere on the patterns for 16 hours, at which time they fixed and processed for immunofluorescence and quantification analyses.

### Immunofluorescence stainings and confocal microscopy

Cells were fixed in 4% paraformaldehyde, permeabilized with 0.3% Triton X-100 in PBS, and blocked in 5%bovine serum albumin (BSA). Samples were subsequently incubated overnight in primary antibody in 1% BSA/0.3% Triton X-100/PBS, followed by washing in PBS and incubation in secondary antibody in 1% BSA/0.3% Triton X-100/PBS. Finally, samples were mounted in Elvanol. The following antibodies were used: *α*-parvin (Cell Signaling; 4026; 1:100), phospho-Myosin Light Chain 2 (Thr18/Ser19) (Cell Signaling; 3674; 1:200), Paxillin (BD Transduction Laboratories; 610051; 1:300). Alexa Fluor 568 and 647 conjugated secondary antibodies (1:300, all from Invitrogen). Actin was labeled with Alexa Fluor 568 (Invitrogen; A12380 1:600), or 647-conjugated phalloidin (Invitrogen; A22287 1:100). All fluorescence images were collected by laser scanning confocal microscopy (SP8X; Leica) with Leica Application Suite software (LAS X version 2.0.0.14332), using 63x oil-immersion objective. Images where acquired at room temperature using sequential scanning of frames of 0.3 *μ*m thick confocal planes (pinhole 1) after which 5 planes encompassing complete cell focal adhesions or actin stress fibers were projected as a maximum intensity confocal stack. Images were collected with the same settings for all samples within an experiment. Quantification of adhesion areas was performed using Fiji (Schindelin et al., 2012). First, ILK, or paxillin focal adhesions were identified using intensity-based thresholding. Subsequently, focal adhesion surface area was measured by Analyze particle tools in Fiji. The cell area (in *μ*m^2^) was measured by manually tracing the cell boundary given by adhesions, phalloidin, and pMLC2 stainigs. The number of focal adhesions were normalized by the corresponding cell area.

### Statistical Analysis

Statistical analyses were performed using GraphPad Prism software (GraphPad, version 8) and Python. Statistical significance was determined by the specific tests indicated in the corresponding figure legends. All experiments presented in the manuscript were repeated at least in 3 independent experiments/biological replicates. No datapoints were excluded from the analyses.

## Supporting information

supplementary figures and legends for supplementary movies

## Acknowledgments

The authors acknowledge Anu M. Luoto for technical assistance, support by the Klaus Tschira Foundation, the state of Baden-Württemberg through bwHPC and the German Research Foundation (DFG) through grant INST 35/1134-1 FUGG (to FG) and Helsinki Institute of Life Science, Sigrid Juselius Foundation, and Academy of Finland (Grant 317597) (to SAW). Molecular graphics and analyses were partially performed with UCSF Chimera, developed by the Resource for Biocomputing, Visualization, and Informatics at the University of California, San Francisco, with support from NIH P41-GM103311.

