## supplementary figures and legends for supplementary movies for "ATP allosterically stabilizes Integrin-linked kinase for efficient force generation"

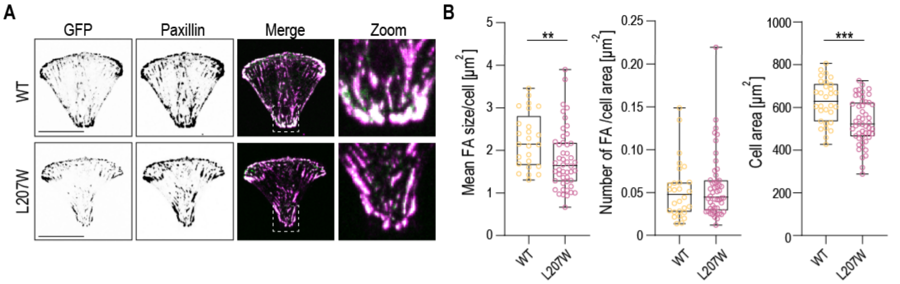


**SFig. 1:** (A)Representative immunofluorescence images of ILK(WT)-GFP and ILK(L207W)-GFP cells stained with paxillin on crossbow micropatterned surfaces. Paxillin localizes at focal adhesions both in ILK(WT)-GFP and ILK(L207W)-GFP cells. (B) Quantification of focal adhesion size/cell, number of focal adhesion/cell and cell area. (n >6 cells/condition/experiment pooled across 4 independent experiments. **p= 0.0052, ***p= 0.0004, Mann-Whitney). Scale bars 20$\mu$m

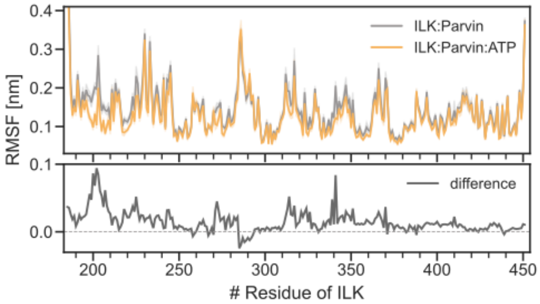


**SFig. 2:** Root mean square fluctuation (RMSF) of ILK holo (orange) and apo (grey) computed for the backbone atoms are shown as a function of residue number. Error bands denote the 95 % confidence interval.

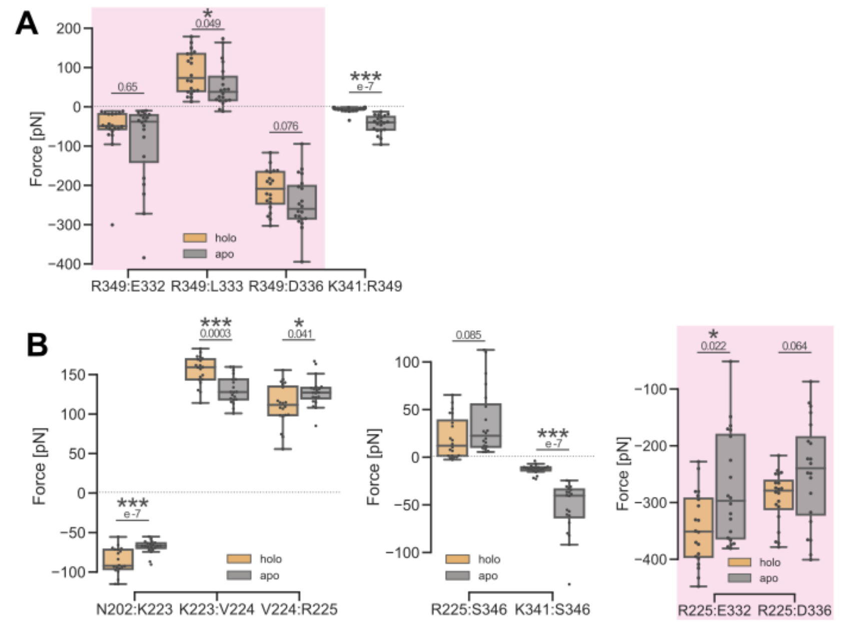


**SFig. 3:** (A,B) Forces from FDA between ILK:ILK and ILK:parvin (pink background) residues for an ATP-dependent pathway of internal force propagation involving ILK R349 and R225, respectively.

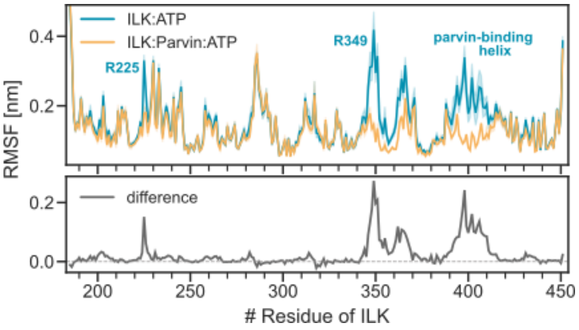


**SFig. 4:** Root mean square fluctuation (RMSF) of ILK in complex with parvin(orange) and without (cyan) computed for the backbone atoms are shown as a function of residue number. Error bands denote the 95 % confidence interval.

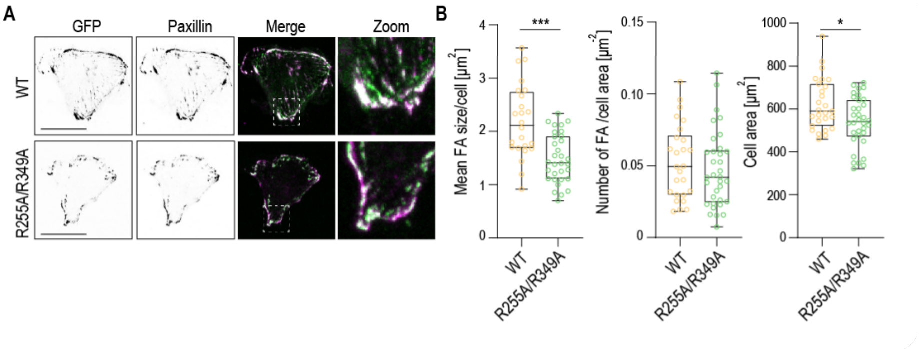


**SFig. 5:** (A)Representative immunofluorescence images of ILK(WT)-GFP and ILK(R255A/R349A)-GFP cells stained with paxillin on crossbow micropatterned surfaces. Paxillin localizes to focal adhesions both in ILK(WT)-GFP and ILK(R255A/R349A)-GFP cells (right panels). (B) Quantification of focal adhesion size and number (n >6 cells/condition/experiment pooled across 4 independent experiments.***p= 0.0001, *p= 0.0260, Mann-Whitney). Scale bars 20$\mu$m.

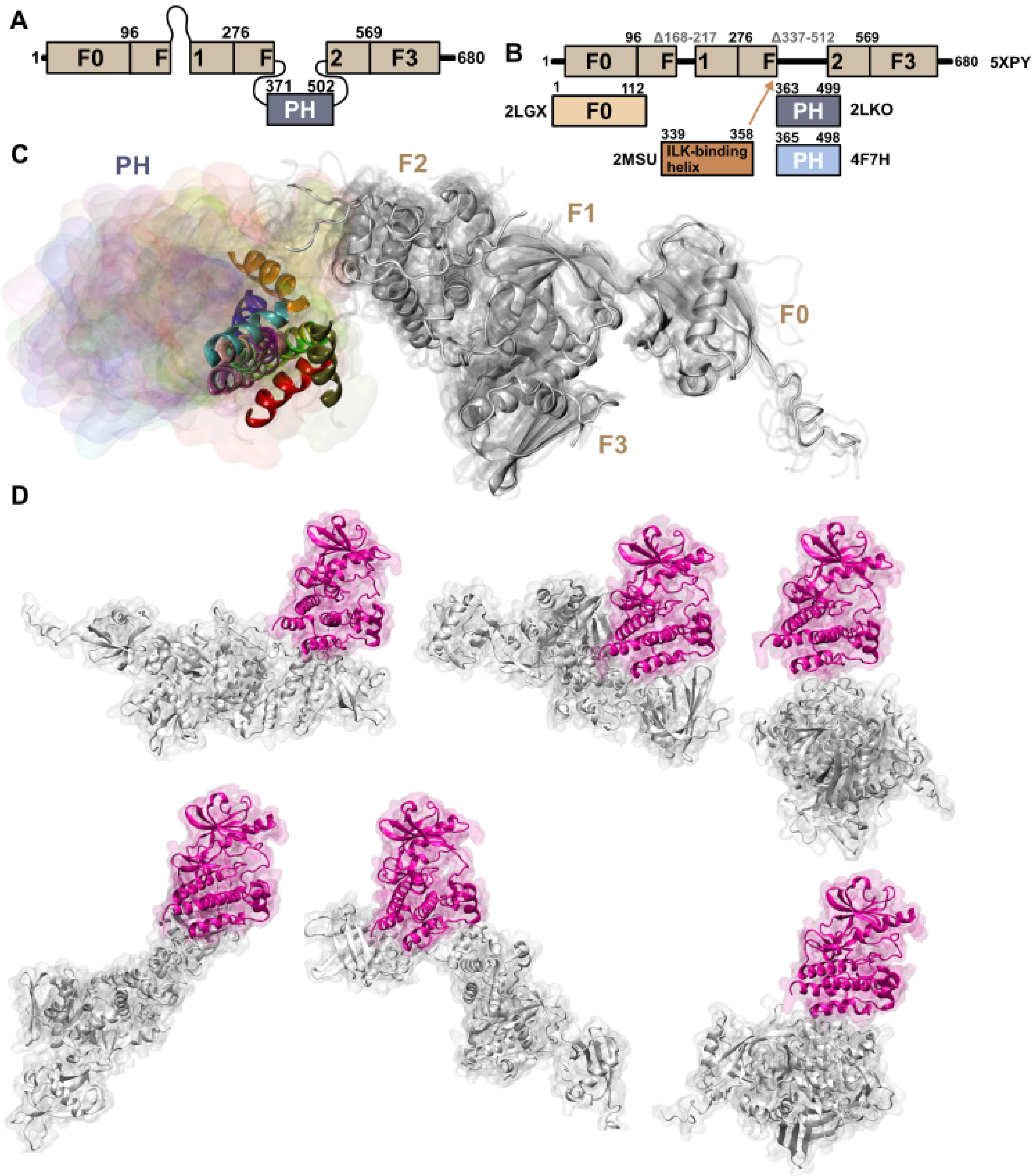


**SFig. 6:** Kindlin-2 modeling and ILK:kindlin-2 docking. (A) Schematic overview of full kindlin-2. (B) Schematic overview of available kindln-2 partial crystal structures with pdb-codes. (C) Full human kindlin-2 homology models with different placements of PH-domains in surface representation in different colors. For visualization purposes one helix within the PH-domain represented in cartoon. (D) 6 exemplaric ILK:kindlin-2 docking poses from guided docking. ILK in pink and kindlin-2 in light grey.

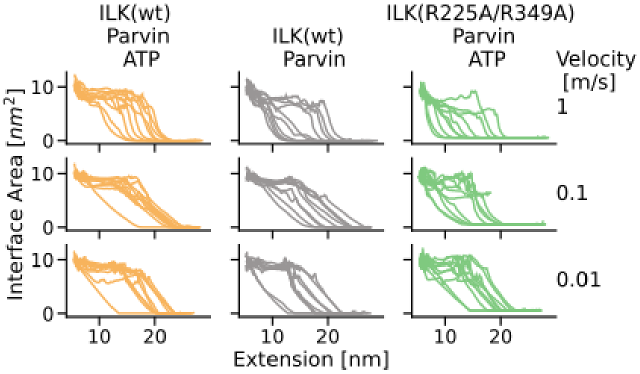


**SFig. 7:** ILK:parvin interface area as a function of extension between the two force application patches for ILK(WT) holo and apo and ILK(R225A/R349A) at three different pulling velocities from $1\,\mathrm{m\,s^{-1}}$ to $0.01\,\mathrm{m\,s^{-1}}$. Trajectories are smoothed with a rolling average.

**SV 1:** Motions captured by PC1. Interpolation between the extreme conformations of apo ILK(wt) along PC1 shown as C-$\alpha$ traces.

**SV 2:** Real time imaging of ILK(WT)-GFP (left) and ILK(L207W)-GFP cells (right) on 8 kPa substrates. Representative movie of ILK(WT)-GFP cells cultured on 8 kPa substrates. Frame rate 1 min/frame. Scale bar 20 $\mu$m.

**SV 3:** Real time imaging of ILK(WT)-GFP (left) and ILK(L207W)-GFP cells (right) on 40 kPa substrates. Representative movie of ILK-WT-GFP cells cultured on 40 kPa substrates. Frame rate 1 min/frame. Scale bar 20 $\mu$m.

**SV 4:** Real time imaging of ILK(WT)GFP (left) and ILK(R255A/R349A)-GFP cells (right) on 8 kPa substrates. Frame rate 1 min/frame. Scale bar 20 $\mu$m.

**SV 5:** Real time imaging of ILK(WT)-GFP (left) and ILK(R255A/R349A)-GFP cells (right) on 40 kPa substrates. Frame rate 1 min/frame. Scale bar 20 $\mu$m.

**SV 6:** Movie of holo ILK(pink) and parvin (cyan) unfolding and complex dissociation in force-probe MD

**SV 7:** Movie of apo ILK(pink) and parvin (cyan) unfolding and complex dissociation in force-probe MD

**SV 8:** Movie of ILK(R225A/R349A)(pink) and parvin (cyan) unfolding and complex dissociation in force-probe MD

**SV 9:** Real time imaging of ILK-/-GFP cell migration. Representative movie of ILK-/-GFP cells cultured on glass and captured by spinning disc time-lapse microscopy at initial (0 h) and 6.30 h time. Frame rate 30 min/frame. Scale bar 20 $\mu$m.

**SV 10:** Real time imaging of ILK(WT)-GFP cell migration. Representative movie of ILK(WT)-GFP cells cultured on glass and captured by spinning disc time-lapse microscopy at initial (0 h) and 6.30 h time. Frame rate 30 min/frame. Scale bar 20 $\mu$m.

**SV 11:** Real time imaging of ILK(L207W)-GFP cell migration. Representative movie of ILK(L207W)-GFP cells cultured on glass and captured by spinning disc time-lapse microscopy at initial (0 h) and 6.30 h time. Frame rate 30 min/frame. Scale bar 20 $\mu$m.

**SV 12:** Real time imaging of ILK(R255A/R349A)-GFP cell migration. Representative movie of ILK(R255A/R349A)-GFP cells cultured on glass and captured by spinning disc time-lapse microscopy at initial (0 h) and 6.30 h time. Frame rate 30 min/frame. Scale bar 20 $\mu$m.
